# Multilayer view of pathogenic SNVs in human interactome through *in-silico* edgetic profiling

**DOI:** 10.1101/299891

**Authors:** Hongzhu Cui, Nan Zhao, Dmitry Korkin

## Abstract

Non-synonymous mutations linked to the complex diseases often have a global impact on a biological system, affecting large biomolecular networks and pathways. However, the magnitude of the mutation-driven effects on the macromolecular network is yet to be fully explored. In this work, we present an systematic multi-level characterization of human mutations associated with genetic disorders by determining their individual and combined interaction-rewiring, “edgetic”, effects on the human interactome. Our *in-silico* analysis highlights the intrinsic differences and important similarities between the pathogenic single nucleotide variants (SNVs) and frameshift mutations. We show that pathogenic SNVs are more likely to cause gene pleiotropy than pathogenic frameshift mutations and are enriched on the protein interaction interfaces. Functional profiling of SNVs indicates widespread disruption of the protein-protein interactions and synergistic effects of SNVs. The coverage of our approach is several times greater than the recently published experimental study and has the minimal overlap with it, while the distributions of determined edgotypes between the two sets of profiled mutations are remarkably similar. Case studies reveal the central role of interaction-disrupting mutations in type 2 diabetes mellitus, and suggest the importance of studying mutations that abnormally strengthen the protein interactions in cancer. With the advancement of next-generation sequencing technology that drives precision medicine, there is an increasing demand in understanding the changes in molecular mechanisms caused by the patient-specific genetic variation. The current and future *in-silico* edgotyping tools present a cheap and fast solution to deal with the rapidly growing datasets of discovered mutations.

## INTRODUCTION

In the past several decades, tremendous efforts and vast resources invested in the quest to understand the molecular mechanisms underlying human genetic disorders resulted in a significant progress towards this goal Next Generation Sequencing (NGS) methods, ranging from the whole-genome sequencing [1] to single-cell transcriptomics [2], have played an instrumental role in this quest, allowing the researchers to investigate genetic determinants in the healthy and disease tissues or cells of the individuals and across populations [3]. These ongoing disease-centric sequencing projects and genome-wide association studies have provided us with an extensive list of genes associated with the diseases and the corresponding mutations. For example, according to the National Cancer Institute-National Human Genome Research Institute (NCI-NHGRI) catalog of published GWAS projects, there are 2,060 publications describing 14,876 SNVs [4]. The dataset includes almost all common and many rare complex diseases, and it is expected that a comprehensive catalog of nearly all human genomic variations, whether deleterious or neutral, will be available soon [5].

At the same time, substantial improvements have been made to utilize high-throughput “-omics” data, including proteomics, metabolomics, transcriptomics, and others, for the purpose of diagnostics and treatment of the complex genetic disorders [3]. One of such high-throughput system-wide technologies—interactomics, whose goal is to characterize the protein-protein interaction (PPI) network functioning in a cell—has become a prominent direction [6–8]. With the release of several large-scale human interactomes [9–11], these complex PPI networks have been utilized to explore the genotype-to-phenotype relationships on the basis that many proteins function by interacting with other proteins, and thus the network-rewiring genetic effects may lead to the disease phenotype [6, 12].

Along with the development of the NGS-driven applications studying human diseases, many genetic variation and genotype–phenotype databases have been developed to assist scientists to better understand the intricacy of the data [3]. For example, ClinVar [13], a freely accessible public collection of reports of the relationships between the genetic variants and phenotypes, also serves as a central archive for causality predictions, with five standard categories ranging from benign to pathogenic. In addition, a plethora of functional annotations tools for genetic variations have been developed to evaluate the functional effect of genetic variations [3], including our recently developed SNP-IN tool [14]. SNP-IN tool predicts the effects of non-synonymous single nucleotide polymorphisms (nsSNPs) on the corresponding protein-protein interaction, given the known structure or the homology model of the PPI complex. The accurate and balanced performance of SNP-IN tool makes it useful for functional annotation of disease-associated nsSNPs.

Together, the above findings bring us one step closer towards mechanistic understanding of the complex genetic disease. However, it has rarely been possible to translate such a massive amount of information on mutations and their associations with disease into biological or therapeutic insights, and the mechanisms underlying genotype-phenotype relationships remain partially explained [15]. The complex genotype-phenotype relationships among diseases are much more complex than was previously expected. For example, the same gene can be associated with multiple disorders (gene pleiotropy). The traditional “one gene/one enzyme/one function” concept assuming a simple, direct, and linear connection between the genotype of an organism and its phenotype often no longer holds [16]. First, one of the most important questions of how the disease-associated mutations impair protein activities in the context of biological networks remains primarily unclear. Although a few alleles have been well-characterized, the functional information is unknown for over 100,000 disease-associated variants in ClinVar [13]. However, profiling several thousand missense mutations in the experimental lab remains costly and laborious. Armed with our recently developed SNP-IN tool, we could bypass the bottleneck and systematically characterize the genetic mutations faster and at a much lower cost.

Recently, a concept of “edgotype” has been proposed [17], which determines the functional outcomes of genetic variants on a PPI, leading to a new study of edgetic profiling to interpret genotype-to-phenotype relationships using the interactome. Another recent work integrated structural annotation with the interaction data from the main pathway repositories showing that the PPI structural information may play a key role in understanding the biological mechanism underlying the disease process and would be an essential component for the next generation of drug development strategies. Also, it is intriguing to study the effect of the removal of a set of disease-associated proteins and the PPIs they are involved in on the performance of the whole system. A considerable amount of efforts has been devoted in other fields to understanding how the network structure changes as it is degraded through the removal of nodes or edges [18, 19]. The removal of nodes or edges has been mostly based on some global or local measures of their possible importance. However, in a biological system, these components with “great importance” do not always overlap with the disease genes. Thus we expect that the previous studies cannot characterize the network rewiring behavior observed in the network centered around the genes associated with a disease, without determining the effects of genetic variation on the protein stability and protein-protein interaction.

Here, we treat the molecular network as a framework, using which one can integrate different kinds of data, including interaction data, structure data, and functional data, rather than a classical graph structure. Specifically, we consider the most common type of genetic variation, SNVs, and seek to understand how SNVs perturb the network, and how this information can be used in linking the genotype with disease phenotype. We map the deleterious SNPs onto the PPI network using SNP-IN tool and evaluate the widespread edgetic perturbations in the human interactome caused by the pathogenic SNVs. The annotation results also provide an additional layer of information for studying network and lead us to the analysis of cumulative network damage caused by the pathogenic SNPs associated with the same disease. Topological analysis suggests that, compared to the frameshift mutations, SNVs are more likely to be the cause of gene pleiotropy, and both the network centrality and mutation enrichment on the protein interfaces contribute to the pleiotropic effect. The functional annotation results also reveal the high percentage of the pathogenic SNVs that could cause rewiring of the corresponding PPIs and that often work synergistically, to a greater extent than was previously expected. Moreover, the analysis of the cumulative damage induced by the pathogenic missense mutations, based on the recently proposed edge-based robustness measure, shows that this measure is more suitable to characterizing the network rewiring behavior than a node-based measure, and that the disease nsSNVs are more efficient in damaging the network, compared to the random attacks.

When applying this approach to study a disease network centered around the genes associated with Type 2 Diabetes Mellitus (T2DM), we have identified systems enriched with disruptive mutations and determined that the interaction perturbations may span the entire network. We have also discovered that the beneficial mutations often played critical roles in the disease progress. Finally, the clinical data analysis indicates that the cancer patients carrying disruptive mutations in the cancer driver genes correlate are linked to the decreased patient relapse time and survival time, suggesting that the disruptive somatic mutations in cancer patients could be utilized as biomarkers predicting the patient’s prognosis.

## RESULTS

The system-wide edgetic characterization of the pathogenic mutations in the human interactome in this work can be broken down to several stages (Fig. 1). First, we studied the topological properties of the pathogenic non-synonymous single nucleotide variants (nsSNVs) in the network. The main goal of this stage was to determine whether one could differentiate between the different types of human genetic variations, either in terms of clinical importance, *i.e.*, pathogenic nsSNVs versus non-pathogenic nsSNVs, or in terms of structural properties, *i.e.*, pathogenic nsSNVs versus pathogenic frameshift mutations. Next, we wanted to check which of the two most common genetic variations was more likely to be the cause of gene pleiotropy and whether perturbations of specific PPIs caused by these variations play a role in gene pleiotropy. Second, we examined these mutations in a structurally resolved PPI network, INstruct [20]. Specifically, we leveraged structural information on PPI complexes and utilized our SNP-IN tool to annotate the edgetic effects of nsSNVs at the systems level and evaluate the widespread perturbations of pathogenic SNVs in the human interactome, thus adding another layer of functional information in this work. Further, we adopted a concept of network robustness from the field of physics in order to quantify the overall, or cumulative, damage induced by the disease-associated mutations and to correctly characterize the network rewiring behavior. Finally, we collected cancer patients’ clinical data, with the goal to link the network damage caused by mutations with the clinical outcome.

**Figure 1.**
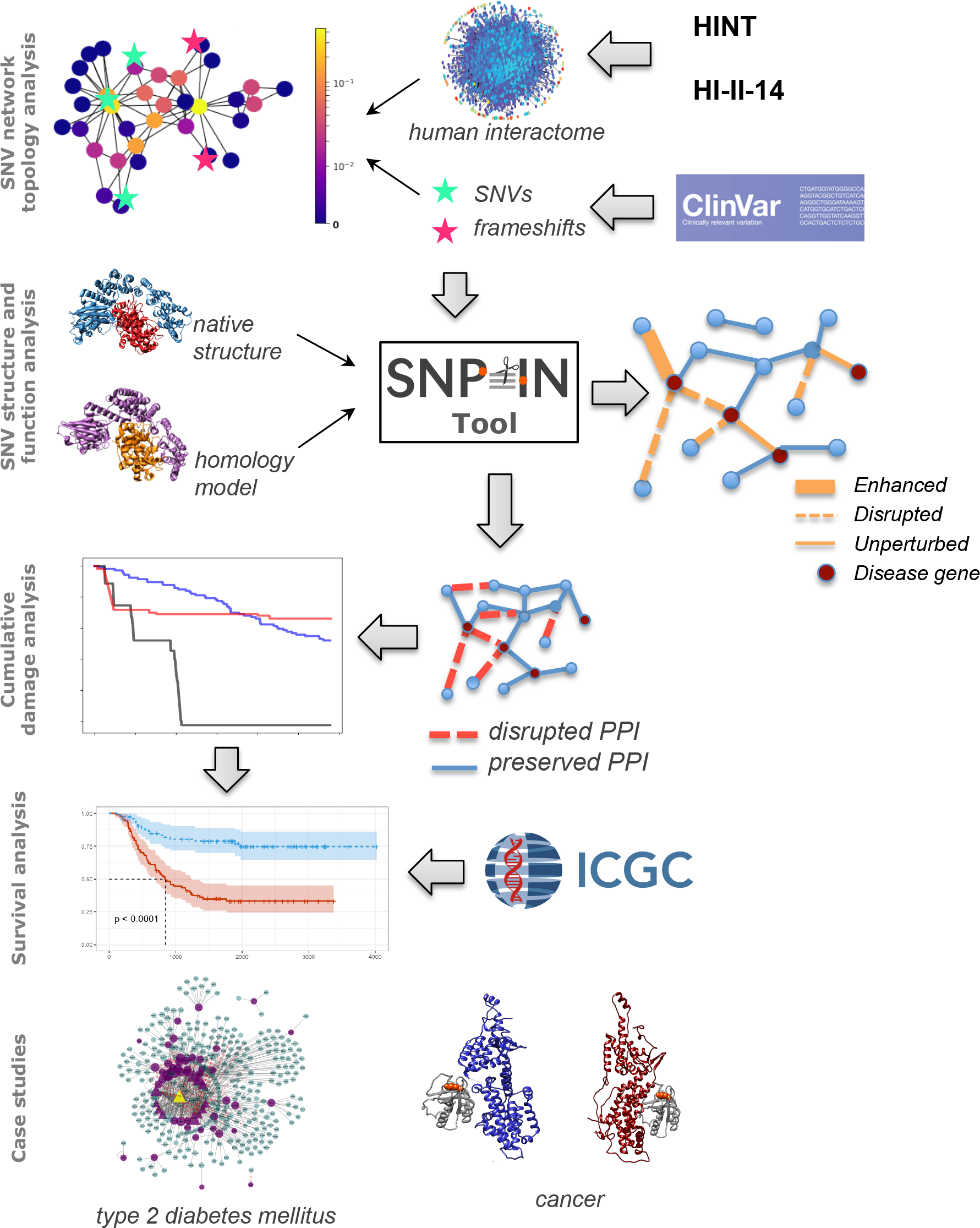
Overview of our approach. First, the topological properties of the non-pathogenic SNVs as well as pathogenic SNVs and frameshift mutations are compared for two interactomes, HINT and HI-II-14. Second, the edgetic profiling of disease SNVs is obtained by applying our SNP-IN tool. Third, the cumulative network damage is studied by applying the principles of the network robustness theory. Fourth, the survival analysis is performed to understand the role of mutations that affect PPI. Last, two large-scale case studies are considered.

For our analysis, two different human interactomes were used and their results compared: HINT [21] and the human interactome project (HI-II-14) network [9]. HINT is a database of high-quality PPIs for human. It integrates several databases and filters out low-quality and erroneous interactions. The second network, HI-II-14, is a reference human protein-protein interactome mapped using a primary yeast two-hybrid array followed by orthogonal validation. The networks present complementary information: the interaction overlap between HINT network (45,226 PPIs), and HI-II-14 network (32,465 PPIs) is only 13,223 PPIs. Next, ClinVar database [13] was used as a source for genetic variants associated with the diseases. ClinVar is a public database where each genetic variant is annotated with some clinical significance for the reported conditions (see *Methods* section for more details). It contains both germline and somatic variants of different types, sizes, or locations. In this work, we curated 11,487 disease nsSNVs distributed across 2,240 genes, 2,719 non-pathogenic nsSNVs in 807 genes, and 6,498 pathogenic frameshift mutations in 1,537 genes. We note that the same gene can carry mutations of different types: for instance, there are 1,039 genes carrying both pathogenic SNVs and frameshifts and 388 genes carrying both pathogenic and non-pathogenic SNVs.

### Pathogenic SNVs share similar centrality properties as frameshift mutations, but are more likely to cause gene pleiotropy

We examined the topological properties of the genes carrying pathogenic SNVs in the two human interactomes by comparing these properties with (i) genes carrying pathogenic frameshift mutations, and (ii) genes carrying non-pathogenic SNVs. SNVs and frameshifts are the two most common genetic variations associated with human disorders [22], so it is natural to study them first in the context of network topology. We evaluated three basic topological properties: node degrees, betweenness, and closeness (Fig 2A) for the HINT interactome and HI-14 interactome separately. The average node degree, betweenness, and closeness for the proteins carrying pathogenic SNVs were 8.6, 1.1 × 10^−4^, and 0.23 in HINT, and 8.3, 1.3 × 10^−4^ and 0.26 in HI-14 interactome. When comparing with the pathogenic frameshift mutations, we found the values for all three properties to be similar (Fig. 2B, Supplementary Fig. S1).

**Figure 2.**
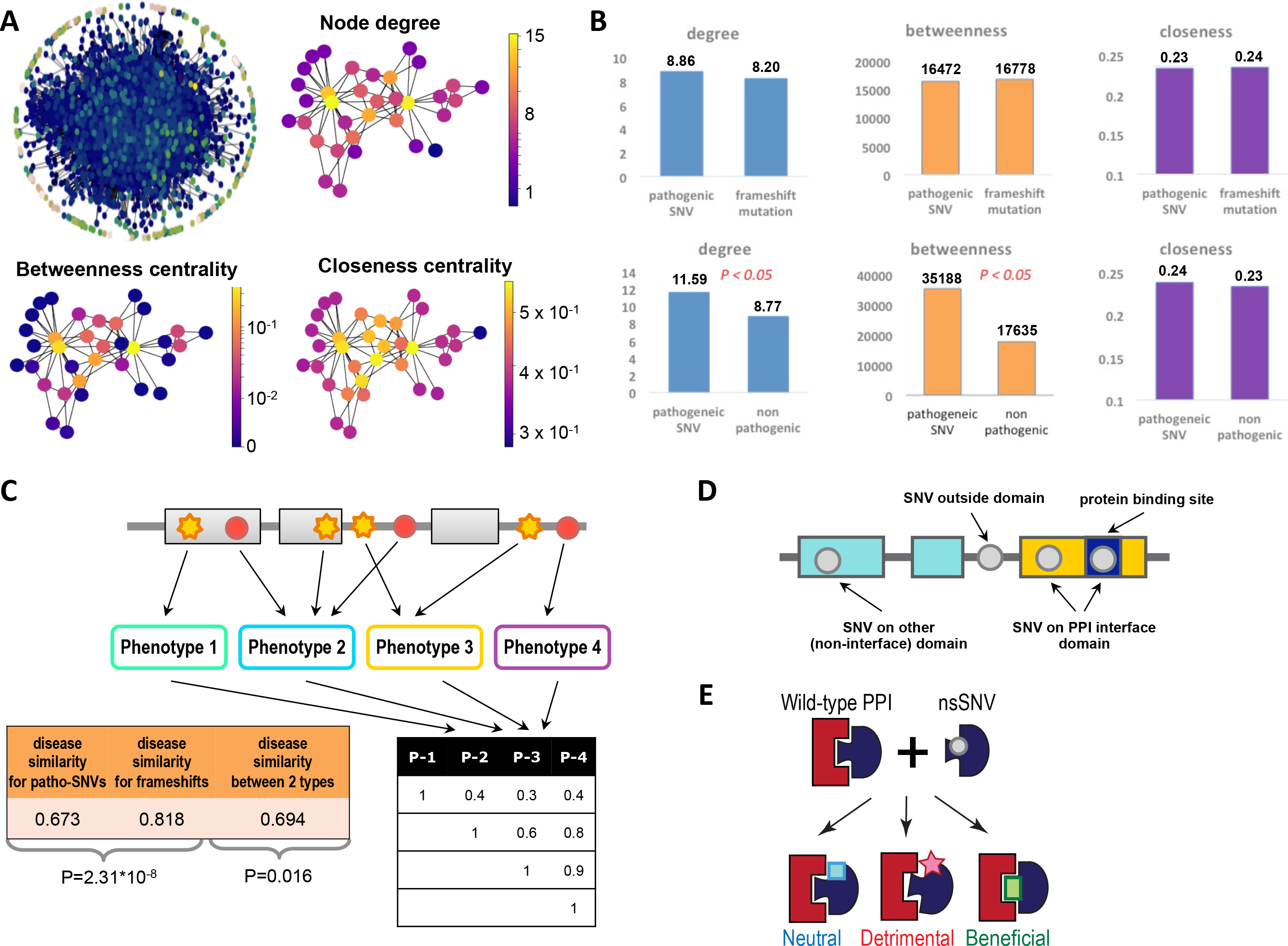
Topological analysis and structural analysis of the pathogenic mutations in the human interactome. (A) Three basic topological characteristics of mutations in the network are calculated: node degree, betweenness centrality, and closeness centrality. Shown are examples of the three measures calculated for the same small network, and the overall scale-free topology of HINT interactome. (B) Comparison of centralities between two groups of mutations calculated for HINT network: pathogenic SNVs vs. pathogenic frameshift mutations, and pathogenic SNVs vs. non-pathogenic SNVs. (C) Basic principles of the analysis of the relationship between gene pleiotropy and mutation source. Red circles represent pathogenic SNVs, yellow stars represent the frameshift mutations. Both mutation types can be associated with different disease phenotypes. For each pair of phenotypes, their similarity is calculated based on the number of common genes associated with both phenotypes. (D) Basic principles of the SNV structural analysis with respect to the protein domain architecture. Grey circles represent SNVs and their location on the protein, while rectangles correspond to the protein domains. (E) Three basic classes of SNVs annotated by SNP-IN tool.

Intriguingly, the comparison of the network properties between the genes carrying pathogenic and non-pathogenic SNVs (Fig. 2B, Suppl. Fig. S2) revealed that the former had significantly higher node degree in HINT interactome (P-value is 0.046, Wilcoxon test) and significantly higher betweenness in both interactomes (P-values are 0.01 and 0.043 for HINT and HI-14 interactomes, respectively, Wilcoxon test). We therefore concluded that the genes with pathogenic SNVs are more central in the network, compared to the genes carrying non-pathogenic SNVs and the changes in the former group of genes are likely to have greater impact on the interactome than similar changes in the latter group. However, the obtained results also implied that these plain topological properties could not differentiate the pathogenic SNVs from pathogenic frameshift mutations. Thus, we next investigated if one could differentiate these two groups of pathogenic variations in a structurally resolved interactome.

Pathogenic mutations are believed to cause the disease in multiple ways [23]. While a frameshift mutation often results in an incomplete protein fragment that is likely to be unfolded or misfolded and thus degraded by the proteosome, a pathogenic nsSNV is likely to produce a full-length protein with a local defect. However, the question of whether such different structural effects on a protein can result in similar effects on a PPI mediated by this protein, the function carried out by this interaction, and a result, the phenotypic change caused by the functional changes, remains largely unanswered. To answer this question, we leveraged the concept of disease phenotype similarity score [24]. The score determines if the two disease phenotypes are similar; the higher the score the more similar two phenotypes are. We calculated the disease similarity caused by above two kinds of pathogenic mutations in each gene and found that the average disease similarity between a pair of pathogenic frameshift mutations is significantly higher than that the similarity between an nsSNV and a frameshift mutation (Fig. 2C). Furthermore, the average disease similarity between a pair of pathogenic nsSNVs in a gene was significantly lower compared to that one between a pair of frameshift mutations.

### Pathogenic SNVs are enriched on the interaction interfaces

As we observed above, the network topology itself could help providing only a high-level view on the effects of the pathogenic nsSNVs. In the recent years, several works proposed to complement the topological network with the structural information [23, 25]. Following the same strategy, we compared the two types of pathogenic mutations extracted in this work by mapping them into a structurally resolved network, INstruct [20]. Specifically, we wanted to find whether the pathogenic mutations have a tendency to accumulate on the protein binding site, and thus contributing to the interaction interface, as opposed to the rest of the protein. The interface enrichment of the pathogenic SNVs would indicate that they are likely to cause the disease through rewiring the corresponding protein-protein interactions (Fig. 2D). The set of disease genes (672 genes) containing at least one pathogenic nsSNV and at least one protein binding site from INstruct was selected to calculate the interface mutation enrichment. We found that the pathogenic nsSNVs were indeed significantly enriched in the protein interaction interfaces: among all the 4,108 disease-associated nsSNVs in the structurally resolved network, 2,781 of them were observed on a PPI interface (Table 1). Furthermore, the pathogenic nsSNVs were found to be under-represented on the other domains not involved in the PPIs (Fig. 2D, Table 1). In contrast, we found that the non-pathogenic SNPs from the same set of the disease genes were not enriched on the PPI interfaces (Table 2). These observations provide strong support for the proposed mechanism causing the disease phenotype through PPI rewiring by the pathogenic SNVs.

**Table 1.** Distribution of SNVs across the protein sequence: 1. pathogenic SNV group; 2. non-pathogenic SNV group. **N** corresponds to the number of SNVs. **OR** corresponds to odds ratio.

|  | Interface Domain |  | Other Domain |  | Non-domain |  |
| --- | --- | --- | --- | --- | --- | --- |
|  | N | OR | N | OR | N | OR |
| 1. | 2,781 | 2.41 | 325 | 0.84 | 1,002 | 0.41 |
| 2. | 180 | 0.70 | 47 | 0.67 | 653 | 1.58 |

**Table 2.**
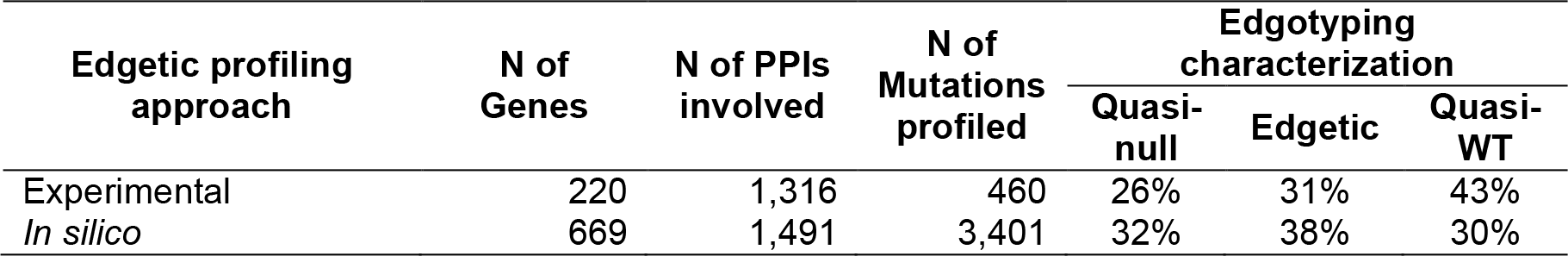
Comparison between the recent large-scale experimental edgetic profiling study [27] and the current study performed using an *in silico* approach [14].

| Edgetic profiling approach | N of Genes | N of PPIs involved | N of Mutations profiled | Edgotyping characterization |  |  |
| --- | --- | --- | --- | --- | --- | --- |
|  |  |  |  | Quasi-null | Edgetic | Quasi-WT |
| Experimental | 220 | 1,316 | 460 | 26% | 31% | 43% |
| *In silico* | 669 | 1,491 | 3,401 | 32% | 38% | 30% |

### Functional annotation of disease SNVs indicates widespread disruption of interactome and synergistic edgetic effects of SNVs

While the enrichment of the pathogenic nsSNVs on the interaction interfaces suggest that pathogenic nsSNVs play an important role in the protein interactions, their mere presence on the PPI interface does not guarantee that each such mutation would have a functional effect on the interaction. Our recently developed SNP-IN tool was designed to differentiate between the interaction-neutral nsSNVs and those ones affecting the interaction [14]. SNP-IN tool accurately predicts the effects of non-synonymous SNVs on PPIs, given the interaction’s structure. It is designed as a set of classifiers leveraging a new Random Forest self-learning protocol. A 3-class classification problem was considered in our study (Fig. 2E), where the classes corresponded to the three functional effects of SNVs on the protein interaction assigned based on the difference between the binding free energies of the mutant and wild-type complexes: detrimental, neutral, and beneficial (see Methods for the definitions).

SNP-IN tool requires the structure of the PPI complex, which comes from either an experimentally resolved structure or an accurate model from the homology modeling approach. In total, we were able to provide at least partial structural characterization of 1,491 PPIs. To meet this requirement, on one hand, we performed a comprehensive search in the PDB database [26] (see *Methods*), obtaining 499 experimental structures with structurally resolved PPI interface. Furthermore, we have obtained the full-length homology models for 818 PPIs and partial domain-domain interaction homology models for 174 PPIs. There has not been a common agreement on to what extent the disease associated mutations could affect the PPIs. One study concluded that only a small number of disease-associated mutations were expected to specifically affect PPIs. However, it has been suggested that perturbations of PPIs (disruptions or enhancements) played an important role in the pathogenesis of many disease genes, more than previously expected [23]. Our results showed that, among all 3,401 SNVs annotated by SNP-IN tool, which accounted for about 1/3 of the total disease-associated SNVs we collected, 2,592 SNVs (76.2%) were predicted as detrimental to at least one PPI that the corresponding disease protein was involved in, and 48 SNVs (1.4%) were labeled as beneficial.

Further, we explored whether these pathogenic mutations tend to work synergistically or antagonistically. We grouped the beneficial and neutral mutations into a new class, labeled as *interaction preserving*, and named the detrimental mutations as the second class, *interaction disrupting* (Fig 3A). We then defined a synergistic genetic interaction as a mutation pair that had the same effect for the corresponding PPI, either preserving or disruptive. Similarly, we defined the antagonistic interaction as a mutation pair that had the opposite PPI rewiring effect according to the SNP-IN tool annotation. Based on this definition, we focused on the PPIs with at least two annotated nsSNVs. These mutations could be on the same protein or located on the two separate interacting partners. However, they should target the same interaction. In total, we collected 1,491 PPIs with at least two nsSNVs. We found that 24,922 mutation pairs have the same disruptive effect on the same protein-protein interaction, while 9,334 mutation pairs had the same interaction preserving effect, which accounts for 55.1% and 20.6% respectively in the all the 45,205 possible pairwise mutation combinations within the same interaction. At the same time, we had 10,949 antagonistic mutation pairs, accounting for 24.2%. Further, we excluded the mutations with the neutral effects in this analysis and focused the PPIs containing beneficial mutations. We found that only 506 pairs of mutations tend to work antagonistically, a small percentage of all possible mutation combination (11,859). These results suggest that genetic mutations tend to work synergistically and can be explained by the fact that an individual mutation might not be “disruptive” enough to cause a major dysfunction of the corresponding protein-protein interaction, while a group of two or more mutations with the same rewiring effect could be sufficient.

**Figure 3.**
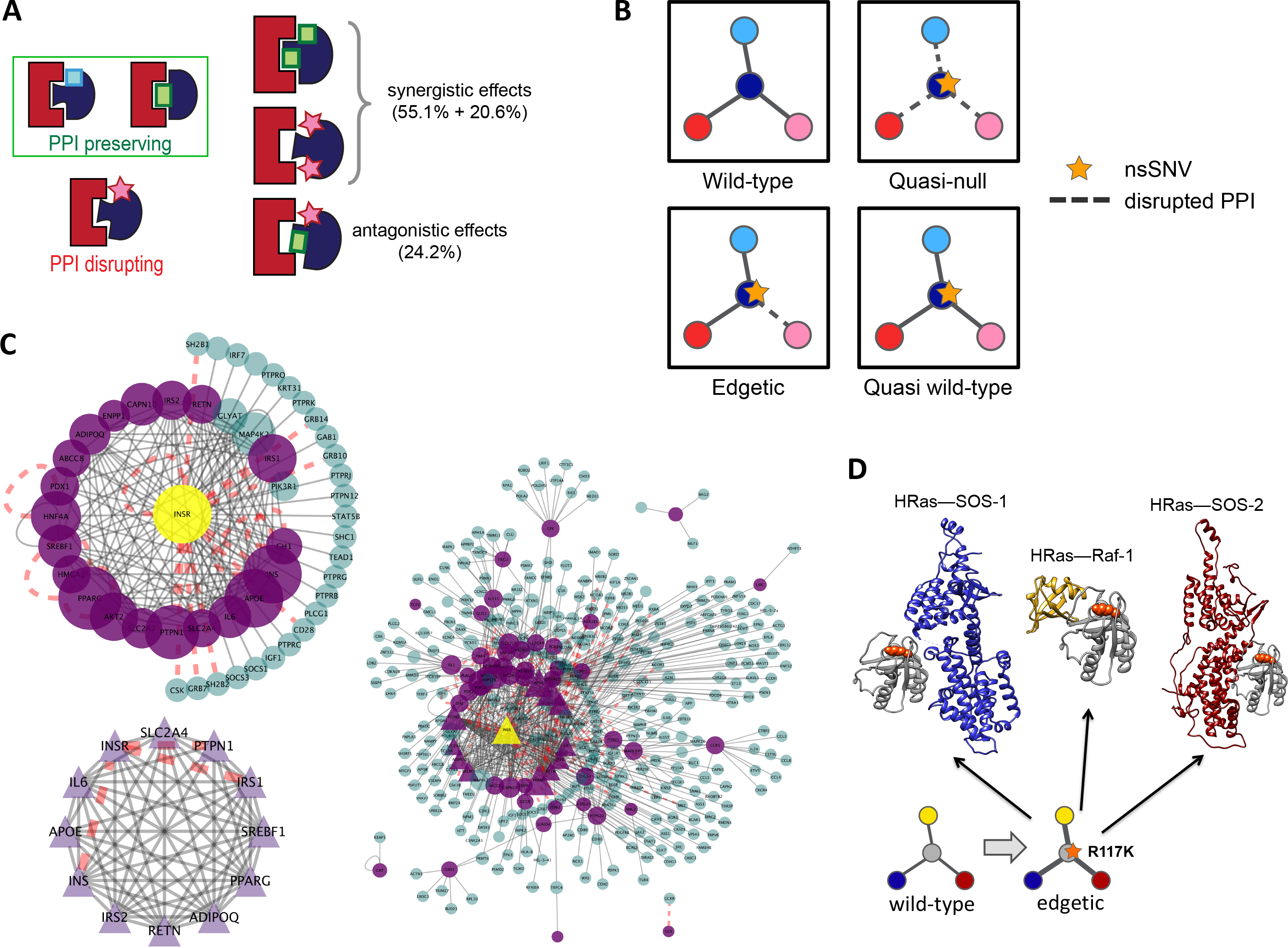
Analysis of the functional annotation results by SNP-IN tool and its application to two case studies. (A) Using the basic classes of SNVs from Fig. 2E, two basic classes of network perturbing mutations are defined: interaction preserving and interaction disrupting. Shown on the right-hand side is the basic principle of the antagonistic and synergistic effects of mutations. (B). Basic edgotypes used in this work. The first one is the wild type interactions. The other three are showing different effects of SNVs on PPI: quasi-null, edgetic, and quasi-wildtype. (C) Case study of the T2DM-centered network. Shown on the right-hand side is the visualization of the entire T2DM-centered network. The central yellow triangle corresponds to INSR gene, which is surrounded by its interaction partners. The purple nodes correspond to the diabetes genes. The triangle nodes correspond to the genes involved in the clique subnetwork. The size of the node corresponds to the node degree. The red dash lines are the disrupted interactions. The left top network is a subnetwork that focuses on INSR gene and its interacting partners. The left bottom subnetwork corresponds to a clique of diabetes genes identified in the network. (D) Case study of HRAS gene and the beneficial mutation on it. An nsSNV, rs104894227, located on HRAS gene is predicted to enhance three independent PPIs: HRas and Raf-1, HRas and SOS-1, as well as HRas and SOS-2 The enhancement of PPIs caused by rs104894227 could explain possible malfunction of the molecular mechanisms underlying the interactions.

### Comparison with experimental edgetic profiling shows greater prediction coverage of the in-silico approach

Recently, the first large-scale edgetic profiling of the missense mutations associated with disease phenotypes has been done by using interaction assays [27]. Specifically, 2,449 mutant proteins and their 1,072 corresponding WT proteins were screened against all partners found in the human interactome HI-II-14 [9]. In total, the interaction profiles for 460 mutant proteins and their 220 WT counterparts were obtained resulting in 521 perturbed interactions found in 1,316 PPIs. The work also provided systematic measurements of the PPI profile changes caused by mutations using a strategy referred to as “edgotyping” [17]. The effects of missense disease mutations on PPIs were grouped into several major categories (Fig 3B): mutations causing no apparent detectable change in interactions (“quasi-WT”), mutations causing specific loss of one or several interaction (“edgetic”), and mutations causing a complete loss of all interactions (“quasi-null”).

Here, we wanted to compare our *in-silico* edgetic profiling method with the experimental profiling approach in order to find the advantages or disadvantages of the former. We found that our computational approach has a significantly higher coverage in terms of number of genes, mutations, and PPIs being profiled (Table 2): in total, we have systematically characterized the effects of 3,401 mutations carried by 669 proteins on 1,491 PPIs. Next, focusing on a missense mutation set for which the corresponding protein has two or more interaction partners, the experimental edgetic profiling identified 26% of the mutations as quasi-null, 31% as edgetic and 43% as quasi-WT. In our case, for a mutation set meeting the same criteria, we determined 32% of them to be quasi-null, 38% as edgetic and 30% as quasi-WT. The distributions of quasi-null, edgetic, and quasi-WT alleles were statistically indistinguishable between the experimental and computationally predicted datasets. Interestingly, in spite of the highly similar distributions, the overlap between the mutation sets from the experimental analysis and our work was minimal: only 56 mutations carried by 33 genes, which is ~4% of a total of 889 genes and 1% of 4,862 mutations considered in both edgetic profiling studies, were shared among the two mutation sets, demonstrating great complementarity between the two approaches.

### Cumulative damage analysis of PPI network reveals network rewiring behavior caused by genetic mutations

Next, we wanted to estimate the synergistic rewiring effect of pathogenic mutations on the whole interactome. To do so, we tested several measures that quantify the cumulative damage caused by a group of mutations. The idea of cumulative damage is exactly opposite to the idea of the network robustness that is commonly used in the network theory to describe the ability of a network to withstand malicious attacks damaging it. First, we performed the node-based cumulative damage analysis on each of the two individual interactomes (Fig 4A, 4B). As a result, we observed similar trends in both interactomes, in spite of the fact that these two interactomes were distinct, overlapping only over a small subset of PPIs and proteins. In particular, we found that both HINT and HI-II-14 interactomes were robust to the “random failure” (*i.e.*, random removal of protein nodes) but vulnerable to “malicious attack” (*i.e.*, the removal of nodes based on the size of the largest network component; nodes whose removal reduces the largest component the most are removed first), which was consistent with the previous findings [15].

**Figure 4.**
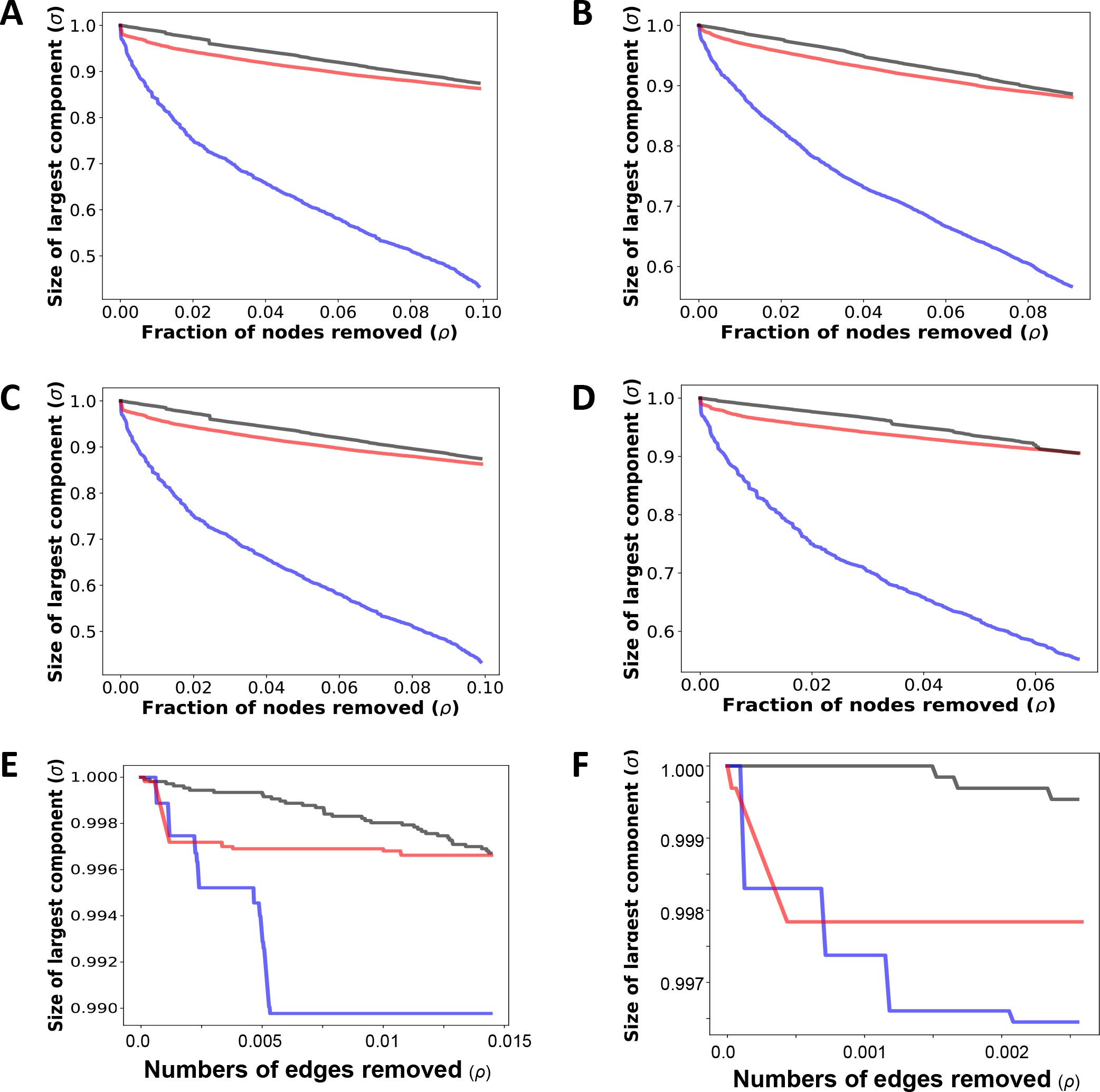
Cumulative damage analysis. Cumulative damage calculated for both interactomes, HINT (panels A, C, and E) and HI-II-14 (panels B, D, and F). Grey lines correspond to the random attack strategy, where the nodes are removed randomly. Blue lines correspond to the removal of nodes based on the size of the largest network component; nodes whose removal reduces the largest component the most are removed first. Red lines correspond to the cumulative damage done exclusively by either pathogenic SNVs or pathogenic frameshift mutations. Three robustness strategies were used and compared: (A), (B) correspond to the cumulative damage calculated using node-based cumulative damage measure, with red lines corresponding to the cumulative damage done exclusively by the pathogenic SNVs that rewire the interactome; (C), (D) correspond to the cumulative damage calculated using node-based cumulative damage measure, with red lines corresponding to the cumulative damage done exclusively by the pathogenic frameshift mutations; (E), (F) correspond to the cumulative damage calculated using edge-based cumulative damage measure, with red lines corresponding to the cumulative damage done exclusively by the pathogenic SNVs disrupting the PPIs.

We also found that when we used the node-based cumulative damage measure to estimate the amount of changes in the interactome due to edgetic effects of the pathogenic SNVs, the network damage caused by the pathogenic SNVs was similar to the random failure (Fig 4A, 4B). However, such insignificant cumulative damage of the network caused by a random failure might not be sufficient to exhibit the disease phenotype, and was contradictory to the idea that disease phenotype could be caused by the network perturbations [17]. Considering the widespread perturbations of PPIs caused by the pathogenic SNVs across a broad spectrum of genetic diseases, it raised a question if the node-based definition of cumulative damage was an accurate measure to characterize the network damage. To substantiate our contention, we also performed the same analysis for the frameshift mutations for both HI-II-14 and HINT interactomes. As discussed earlier, a frameshift mutation often results in an incomplete protein fragment that is typically degraded, which corresponds to a node removal in the interactome. On the other hand, nsSNVs are likely to produce a full-length protein with a potential defect in the corresponding PPI(s), which would correspond to some missing edges but the node associated with the protein is likely to remain intact. Therefore, based on the edgotyping classification, most frameshift mutations would fall into the quasi-null category and are supposed to cause very different rewiring effect on the network. However, our results using the node-based cumulative damage measure (Fig 4C, Fig 4D) suggested that the frameshift mutations have the similar behavior as the pathogenic nsSNVs. This further confirmed the fact that the node-based cumulative damage measure definition could not differentiate the rewiring behavior of nsSNVs and frameshift mutations.

Following the above analysis, we next adopted a more reasonable edge-based definition of cumulative damage. Specifically, we leveraged an edge removal scheme proposed in a recent paper that studied edge-based robustness [19]. We found that in both HINT and HI-II-14 interactomes, the pathogenic mutations could cause more severe damage than a random failure (Figs. 4E, 4F), and the network damage was more similar as the one during malicious attacks (*i.e.*, the removal of edges based on the on the size of the largest network component; edges whose removal reduces the largest component the most are removed first). These results suggested that the edge-based definition was more suitable to characterize the network damage caused by the pathogenic SNVs and might be helpful in assessing phenotypic changes driven by the PPI-rewiring mutations.

### Network analysis identifies a disrupted network clique of proteins associated with type 2 diabetes mellitus

As a first case-study, we carried out the analysis of an interaction network centered around proteins associated with type 2 diabetes mellitus (T2DM). The important role of protein-protein interactions in T2DM was recently proposed [28–30]. Most of these works focused on integrating different sources of data to discover novel candidate genes for T2DM. Here, we aimed at studying the mutation-induced rewiring of the T2DM-centered PPI network. First, we curated 131 T2DM genes from ClinVar database, together with 346 pathogenic SNVs on those genes. To define a T2DM-centered PPI network, we extracted the interaction partners for the T2DM proteins from both HINT and HI-2014 interactomes. Together, we curated 655 interactions. Based on the edgetic profiling done by SNP-IN tool, we were able to annotate 185 T2DM-related mutations, and 139 of them were labeled by SNP-IN tool as disruptive, suggesting global rewiring of the PPI network in T2DM (Supplementary Table S3).

The analysis provided us with several interesting findings. First, we found that T2DM-related genes formed a clique (Fig. 3C). To find how tightly the genes associated with T2DM are connected with each other, we used the MCODE [31] in Cytoscape [32] to perform graph clustering. The highest-scoring cluster consists of 12 genes, which are all associated with T2DM, and 62 PPIs. Based on the SNP-IN tool annotation, we observed 3 disrupted interactions inside a clique (Fig. 3C). Such an inter-connected cluster could be a central functional hub of the T2DM network, thus the PPI perturbations inside the clique could play a central role in the disease phenotype.

Further analysis revealed that among the genes associated with T2DM, *INSR* carried the highest number of disruptive mutations. The *INSR* gene encodes a transmembrane insulin receptor (UniProt ID: P06213), a member of the receptor tyrosine kinase family that mediates the pleiotropic actions of insulin and plays a key role in the regulation of glucose homeostasis [33]. Binding of insulin leads to the phosphorylation of several intracellular substrates, including insulin receptor substrates (IRSs). Two main signaling pathways, *PI3K*-*AKT*/*PKB* and the *Ras*-*MAPK*, are activated following the phosphorylation of IRSs. The *PI3K*-*AKT*/*PKB* pathway is responsible for most of the metabolic actions of insulin, and the *Ras*-*MAPK* pathway regulates specific gene expressions and cooperates with the *PI3K*-*AKT*/*PKB* pathway. *INSR* was found to be involved in the interactions with the majority of known T2DM-associated genes (Fig 3C); it is also a member of the clique subnetwork described above. We annotated 21 mutations associated with *INSR*, and 17 of them were annotated as disruptive. Strikingly, each of the 11 protein-protein interactions that *INSR* participated in, was disrupted by at least one pathogenic SNV, suggesting that, combined, the mutations effectively stop the functioning of this gene.

### Interaction enhancing mutations provide new insights into transient interactions and their roles in diseases

In our second case study we investigated potential roles of a small number of pathogenic nsSNVs that were determined beneficial, *i.e.* causing a significant increase in binding affinity. Recent work [27] suggested that the gain of interactions may be a rare event in human disease. Furthermore, the network topology does not get affected by such mutations, at least when the network dynamics is not considered. Therefore, we hypothesized that the beneficial mutations are mainly involved and affected the transient PPI. A permanent interaction is typically long-term, stable, and irreversible. On the other hand, transient protein complexes form and break down recurrently, with the involved proteins often interacting in a brief period of time and in a reversible manner [34].

We first would like to check how many interactions that the beneficial mutations affected were the transient interactions. Determining the permanent of transitive state of a PPI is an extremely challenging task. Experimental methods that can characterize the transient or permanent state of a PPI are laborious and costly, and a high-throughput characterization of all interactions in the interactome has yet to be carried out. Here, we chose a computational approach, NOXclass [35], which determines the protein-protein interaction type as either obligate or non-obligate. Non-obligate PPIs constitute of proteins that can form stable well-folded structures alone, obligate protein complexes are mainly involved with those proteins that are unstable on their own and become stabilized only through an interaction. Typically, the obligate interactions are permanent, whereas non-obligate interactions are more likely to be transient [36]. Using NOXclass, among all the 55 interactions, we were able to obtain prediction results for 45 interactions. And about 50% are predicted to be non-obligate, suggesting a possible mechanism of action for beneficial mutations—stabilizing the transient complexes.

We further studied an interesting case of one nsSNV, rs104894227, located on *HRAS* gene (protein HRas; Uniprot ID: P01112) that potentially enhanced three independent PPIs: HRas and Raf-1, HRas and SOS-1, as well as HRas and SOS-2 (Fig 3D). *HRAS* gene is a member of the Ras oncogene family encodes a protein located at the inner surface of cell membrane. Mutations in *HRAS* were found to associate with conventional follicular carcinoma [37] and Costello syndrome [38]. The enhancement of PPIs caused by rs104894227 could explain possible malfunction of the molecular mechanisms underlying the interactions. For instance, the transient interaction between HRas and SOS-1 has been known to maintain a delicate balance between being а strong enough connection to facilitate the nucleotide release and being ready to "unzip" for acceptance of the new nucleotides [39]. The PPI-enhancing mutation is likely to break this balance, potentially affecting intracellular signaling pathways that control cell proliferation and differentiation. Another interesting functional impact is in strengthening HRAS-Raf-1 interaction. A counterintuitive phenomenon was described where RAF inhibitors were found to enhance ERK signaling, facilitating tumor cell proliferation, an adverse effect that was seen with RAF inhibitors in melanoma patients [40]. This phenomenon was explained only recently by the fact that the inhibitors promote RAS−RAF association by disrupting RAF kinase domain autoinhibition [41]. The above missense mutation is also likely to play a role in promoting RAS-RAF association, resulting in a similar phenotype. Development of inhibitors for HRas-RAF and HRas-SOS interactions have been discussed as promising directions in cancer therapeutics [39, 41]. The identified edgotypes may be useful in modifying these therapeutic strategies to account for the presence of PPI-enhancing mutants.

### Interaction disrupting mutations on cancer drivers correlate with decreased survival

Lastly, to explore clinical significance of the disruptive mutations, we studied the survival rates and relapsing times of the corresponding cancer patients. Specifically, we found that the acute myeloid leukemia and liver cancer patients with the disruptive somatic mutations carried by the cancer drivers would suffer decreased survival time and relapse time. We first curated a list of 869 high-confidence cancer genes from the Cancer Gene Census [42] and recent literature [43] (see Methods). Among these 869 genes, 227 genes had pathogenic SNVs previously found to be implicated in the cancer progress (Fig 5A). By applying SNP-IN tool, we were able to provide annotation about the effects of these SNVs on PPIs for half of the set, 107 genes. We note that SNP-IN tool did not cover all the mutations for these genes, since some of the mutations could not be mapped to the structures of PPI complexes. In summary, we annotated 784 mutations for these 107 genes, and 58.3% of them showed to be disruptive to PPIs. The top five cancer genes with the highest numbers of disruptive SNVs include *MLH1*, *MSH2*, *STAT3*, *SOS1*, and *VHL* (Fig 5B). On average, a cancer gene carries 18 pathogenic SNVs, and more than 1/3 of them are annotated as disruptive (Fig 5C, Supplementary Table S4). This suggests that mutations in cancers might target protein-protein interactions, and the corresponding rewiring could be a key factor in driving the cancer progress.

**Figure 5.**
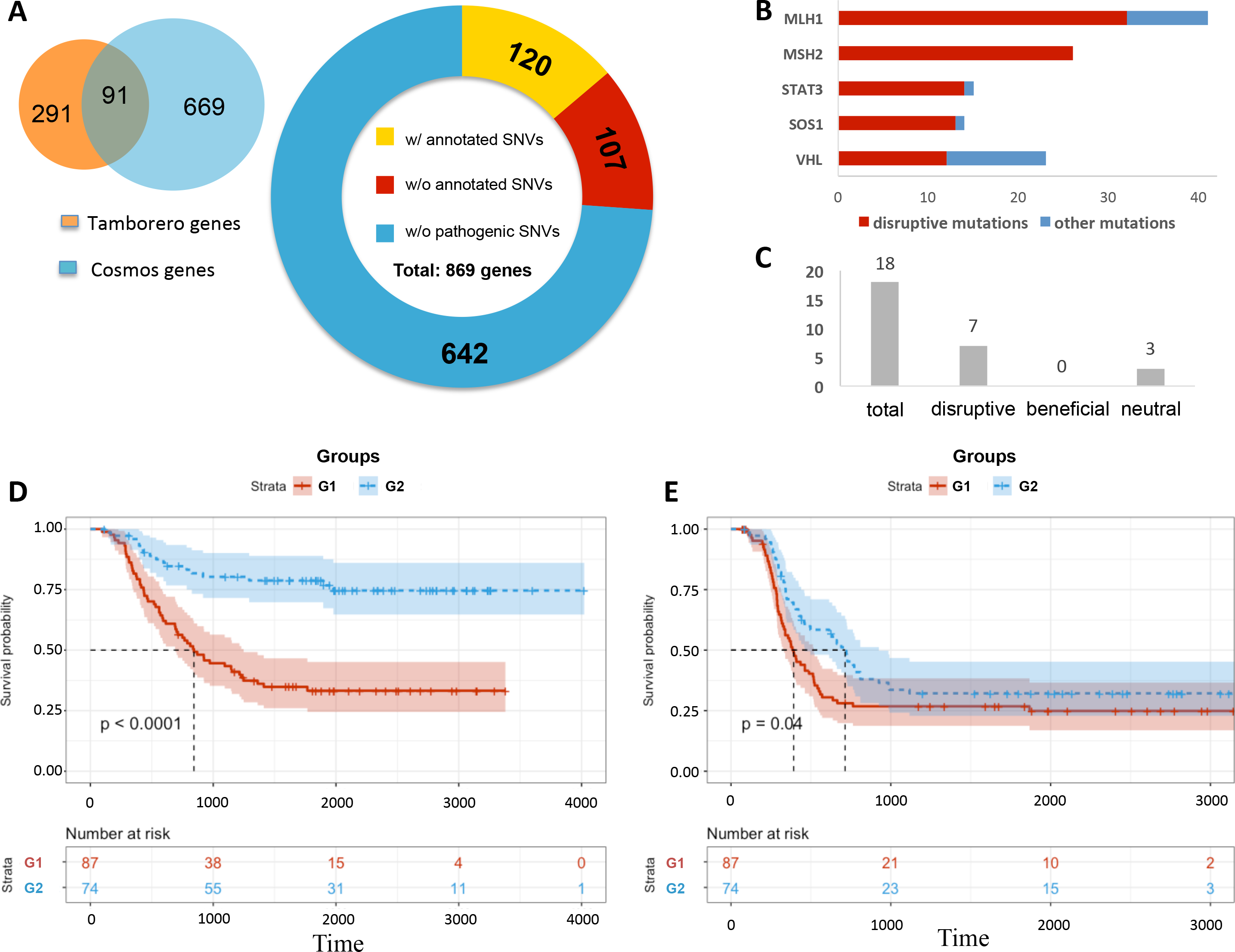
Pathogenic SNVs on the cancer driver genes and their role in the clinical outcomes. (A) Basic statistics of cancer driver genes used in the studies (left) and the annotated SNVs (right). The left part of the panel is showing two main sources of cancer drivers, COSMOS gene consensus and literature collection (Tamborero). (B) Functional annotation of the pathogenic SNVs on cancer drivers. Shown are the top 5 cancer driver genes with the highest numbers of disruptive mutations. The red bar corresponds to disruptive mutations, the blue bar corresponds to other mutations. (C) The average number of mutations of each type on a cancer driver. (D) The survival analysis for the survival time in cancer patients. Based on the edgetic profiling of their mutations, the patients are divided into two groups: G1 (red) includes the patients with the disrupted cancer-centered network, and G2 (blue) corresponds to the patients with the cancer-centered network undisrupted. The lower bar shows the number of patients at risk at the different time points for these two groups. (E) Similar survival analysis for the relapse time. The groups are defined in the same way as in (D).

We next collected the somatic mutations from several cancer sequencing projects and linked the clinical results with the genotyping information of cancer patients with the goal to understand the predictive power of the mutations associated with protein-protein interactions in cancer. The initial idea was to compare the survival statistics between the two groups of cancer patients, one carrying disruptive mutations on the cancer driver genes and another carrying any other mutation (i.e., primarily neutral). The genetic variation data were processed using several bioinformatics tools [44] to curate the corresponding gene information and the mutation position on the protein sequence (see *Methods*). We required that the selected cancer driver genes had the higher mutation rates than the background mutation rate, focusing on nsSNVs from these highly mutated cancer driver genes, and discarding all non-coding variations, nonsense mutations, indels, and other variants. The analysis of the resulting datasets suggested that while subsets of mutations were annotated as neutral and disruptive, some the patient groups were not big enough to carry out the unbiased statistical tests. As a result, we studied a more general question by selecting the groups of patients with mutations located on the PPI interface versus patients with mutations located outside the interface. In this case, the patients were divided into two groups (87/74 for acute myeloid leukemia; 121/113 for liver cancer). Kaplan–Meier statistics was calculated for both groups, and the corresponding statistical significance was calculated using log rank test (see Methods).

We found that the mutations located on the PPI interfaces of the known cancer drivers significantly correlate with the decreased survival and relapse time (Fig 5D, Fig 5E). For acute myeloid leukemia patients, the survival correlation is evident in comparison with those patients with mutations on the corresponding cancer drivers that lie outside the interaction interface (log-rank test p<0.0001 for patient survival time, p = 0.04 for patient relapse time), suggesting that mutations directly associated with protein-protein interactions could be treated as a survival indicator for acute myeloid leukemia patients. A similar correlation between the interaction interface associated mutations and decreased prognosis is seen among the French liver cancer patients (log rank test p<0.0001 for both, patient survival time and relapse time). In sum, the results above demonstrate that mutations directly associated with the protein-protein interactions in the cancer patients present a strong indicator of the patient’s prognosis and the edgetic perturbation of the interactome might play a key role in cancer progress and lead to decreased survival.

## DISCUSSION

In this work, we have presented a systematic multi-layered analysis of the disease interactome using *in silico* edgetic profiling of mutations and leveraging two independent experimentally validated PPI networks. Our high-throughput approach allows drawing the difference between the system-wide distributions and functional effects by the truly damaging pathogenic mutations and mutations that are expected to have minimal or no effect on the protein-protein interactions, while being located on or near the interaction interface. Our analysis of the three basic groups of mutations associated with the diseases, non-pathogenic nsSNVs, pathogenic SNVs, and frameshift mutations, has shown that both pathogenic groups are distinct from the non-pathogenic nsSNVs in their topological distribution on the network. In fact, both pathogenic types of mutations show remarkable similarity in all three network centrality measures, which is surprising given the different nature of the structural, and therefore functional, impact of these two types of mutations. A frameshift mutation typically results in a significant truncation or nonsense-mediated decay, which leads to the loss of all PPIs mediated by this protein [45], while a missense mutation often has a small, localized effect on the protein’s structure, resulting in a loss of a few, and often just one, PPIs [17]. These results suggest that that it is the “topological” role of a protein carrying the pathogenic mutations in the network rather than the type of this mutation that underlies the overall system-wide damaging effect that leads to the disease phenotype.

In spite of their similarity with respect to the network topology, the two types of the pathogenic mutations differ drastically in their capacity to be linked to the pleiotropic effects. A group of detrimental nsSNVs on the same gene is more likely to cause different disease phenotypes than a group of frameshift mutations, which can be explained by the fact that two frameshift mutations are likely to lead to the same deleterious effect, from the point of view of the protein function, with all interactions mediated by this protein being lost. Mutations in a single pleiotropic gene are known to cause different diseases or a wide range of symptoms. However, it is still unknown which type of genetic variation is the main contributing factor to the pleiotropy. Here, based on the edgetic effects analyzed in this work, one could conclude that the disease phenotypes related to the frameshift mutations in the same gene are likely to be more conservative than the disease phenotypes related to the pathogenic nsSNV, and therefore nsSNVs are more likely to be the source of gene pleiotropy. Another distinctive feature of the pathogenic nsSNVs is their enrichment in the PPI interfaces, which is not observed in either non-pathogenic nsSNVs or frameshift mutations, and suggests that many mutations that are associated with the disease phenotypes target the mechanisms behind the macro-molecular interactions

Our recently developed SNP-IN tool has played an important role in systematic characterization of pathogenic SNVs across various human genetic diseases, complementing the recently published large-scale experimental edgetic profiling study [17]. While the accuracy of our, *in-silico*, approach is expected to be somewhat lower, compared to the experimental interaction assays, the coverage of our method is several times higher than the experimental approach: we were able to profile three time as many genes and more than twice as many mutations. Most importantly, when comparing the distributions of the main edgotypes for our prediction with the experimental results, we find the distributions to be nearly identical. Given the minimal overlap of ~4% of shared genes and 1% of shared mutations with the experimental study, the results suggest that our approach can be used to guide future interactomics experiments, suggesting the most promising candidate mutations for the experimental edgetic profiling. Furthermore, SNP-IN tool itself could be improved. While its coverage is currently greater than the experimental approach, our method requires information about the structure of the PPI interface, either from an experimentally resolved structure of macromolecular complex or from an accurate homology model. This structural information is currently limited and was recently estimated to cover ~15% of the human interactome [46]. As a result, only one-third of all pathogenic non-synonymous SNVs extracted from the ClinVar database could be profiled in this work. Developing advanced computational methods that rely on the structures or structural models of the individual interacting proteins instead of protein-protein interactions, while having comparable accuracy, could significantly increase the coverage of an *in-silico* profiling approach.

The edgetic profiling results show the surprisingly widespread perturbations of the human interactome: 76% of disease-associated SNVs are predicted to rewire PPIs. The complex patterns of mutation-induced network rewiring in different diseases lead us to a question if all these mutations incur the comparable amount of damage, and whether the cumulative effect of damaging mutations is amplified through the synergy of their individual effects. We answer this question by developing a cumulative damage analysis that quantifies the mutation-induced network damage using the basic principles of the network robustness theory. Perhaps, the most important finding from this analysis is the fact the traditional node-based measures used in the robustness theory are not well-suited to capture the damage from edgetic effects, and an edge-based alternative should be used instead. We note that the original network robustness concept characterizes how the network withstand failures and perturbation, while our goal is to quantify the amount of damage made by the pathogenic mutations perturbing the network. There are other critical attributes of complex networks that could potentially be included into the definition of cumulative network damage. Future study could consider network efficiency and network navigability. Network efficiency quantifies the exchange of information across the network. Network navigability studies the structural characteristics of many complex networks that support the efficient communication without the global knowledge on their structure. Routing information in networks is a common phenomenon in many complex systems, including biological networks [47]. In the future, it may be helpful to integrate the interactome’s topological structure, edgetic annotation, and navigation strategies to understand how the genetic variations can influence the network efficiency and network navigability.

Finally, we considered several case studies that provide important insights into the mechanisms of complex genetic disorders, such as cancer and type 2 diabetes mellitus (T2DM). While the role of PPIs in diabetes mellitus has been recently recognized, the previous studies could not distinguish between the neutral and network-damaging mutations. Our edgetic profiling of the genes associated with T2DM suggests protein-protein interaction to be the key molecular mechanism that gets malfunctioned in T2DM because of several reasons. First, the majority of the known mutations in T2DM-associated genes are predicted to disrupt the PPIs. Second, PPIs drive the formation of clusters of tightly interconnected T2DM genes. Last, *INSR*, the gene that encodes a transmembrane insulin receptor has been found to carry the highest number of PPI-rewiring nsSNVs, which cumulatively disrupt all twelve interactions in which this protein is involved, disrupting the important metabolic pathways

When studying genes related to cancer, we find a significant proportion of their mutations (more than 58%) to be disruptive. This number could be potentially higher should the coverage of the SNP-IN tool increase. Of special interest is the analysis of beneficial mutations, which strengthen the PPI instead of disrupting it. While the number of annotated beneficial mutations is much smaller compared to the disruptive mutations, the former group is likely to play an important role in disease mechanisms targeting transient interactions. The example of the cancer-related HRAS gene and its interactions that was considered in this work lead to an important conclusion: the knowledge about the PPI enhancing mutations may affect the therapeutic strategies designed to inhibit specific interactions. The last question one can investigate is the importance of PPI-associated mutation in the analysis of the clinical data. Specifically we ask if the knowledge of the fact that the pathogenic nsSNVs are located in the interaction interface can be helpful in predicting the survival rates and relapsing times. Our analysis has shown that the pathogenic mutations on the cancer drivers correlate well with the decreased survival in cancer patients. In the future, the use of an edgetic profiling method with the higher coverage is expected to provide further evidence to the determined correlation.

In conclusion, our *in-silico* edgetic profiling approach aims to provide mechanistic insights into genotype-phenotype relationship. The role of a fast and inexpensive computational edgotyping approach is becoming increasingly important with the rapid growth of the personalized genomics data and ever-increasing catalog of disease-associated variants. Such an approach can also reduce the cost of the experimental interaction assays by prioritizing the genes and mutations according to the predicted edgetic effects.

## METHODS

### A dataset of disease genes and genetic mutations

We collect a list of disease genes and their pathogenic non-synonymous SNVs from ClinVar database [13]. ClinVar is a freely accessible repository of published evidence on the relationships between specific variants and the corresponding phenotypes in human. Specifically, the *clinvar_00-latest.vcf* file is used, followed by data preprocessing, to get a paired gene and mutation list. All mutations collected by ClinVar are grouped into five clinical significance categories, including *Benign*, *Likely benign*, *Uncertain significance*, *Likely pathogenic*, and *Pathogenic*, following the guidance by The American College of Medical Genetics and Genomics (ACMG) [48]. The nsSNVs that belong to categories *Likely pathogenic* or *Pathogenic* are defined as pathogenic for this work, while nsSNVs that belong to the remaining three categories are defined as non-pathogenic. In addition, we extracted from SNV another group of genetic variants, the pathogenic frameshift mutations.

### Extraction of PPI data and construction of PPI network

For the disease network analysis, two human PPI networks are used: HINT network [21] and the experimental human interactome project network (HI-II-14) [9]. HINT (http://hint.yulab.org) is organized as a centralized database of high-quality human PPIs collected from several databases and annotated using both, an automated protocol and manual curation. The human interactome project is another recently released PPI source. It includes a set of binary PPIs that were constructed through by systematically interrogating all pairwise combinations of predicted gene products using yeast-two-hybrid experiments. As a result, HINT and HI-II-14 represent two distinct networks with a small overlap. Instead of merging two networks, we analyze HINT and HI-II-14 networks independently, treating them as the complementary rather than competing views of the human interactome. We expect that different groups of false positive and false negative PPIs exist for each network.

### Topological Analysis of pathogenic SNVs in human interactome

In our analysis of the pathogenic SNVs, we first investigate their topological importance, that is, whether these mutations are located on the proteins that occupy critical positions in the human interactome. To do so, for each interactome we calculate and examine the centrality measures associated with the proteins that carry those SNVs. Specifically, we investigate three major centrality measures in the graph theory: node degree, betweenness and closeness. Previous works suggest that the PPI network topology could encode information about how molecular interactions contribute to the disease phenotypes [49]. These centrality measures are useful to explore the shared properties of genetic architectures underlying genetic diseases.

The simplest measure of centrality in a network is the *node degree*. For a protein in an interactome, the node degree specifies the number of direct interaction partners this protein has. *Betweenness* is another global centrality measure, which determines the number of shortest paths that connect any pair of nodes in the network and also pass through a given node. Formally, the betweenness centrality measure, *C_B_*(*v*), of a vertex *v* is defined as:

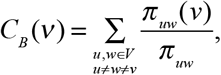

where *π*_*uw*_ is the number of shortest paths between vertices *u* and *w*, and *w*, *π*_uw_(*v*) is the number of such shortest paths that come through vertex *v.* Thus, nodes that occur on many shortest paths connecting pairs of nodes have the higher betweenness.

Closeness centrality provides a rather different view of centrality compared to the above two measures, because it is based on the mean distance between a given node and all other nodes in the network. It is defined as the reciprocal average distance to every other node:

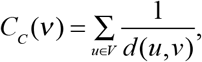

where *d*(*u,v*) is a graph-based distance between the selected node *v* and any other node in the network, *u.* A node with high closeness centrality is, on average, close to the other nodes when using the graph distance. The calculation of the centrality is done using the python Networkx package [50]. To check whether the pathogenic SNVs have higher centrality compared against frameshift mutations, we formulate this question as a statistical test, and apply Wilcoxon test for this task since no prior information about the underlying distribution is known.

### Linking pathogenic SNVs to gene pleiotropy

Pathogenic SNVs and pathogenic frameshift mutations have been suggested to affect the phenotype in different ways [23]. A missense mutation is likely to affect one or several specific interactions. On contrary, a frameshift mutation often causes the loss of all the interactions in which the mutated protein is involved. The mutation-induced perturbations of the network properties give rise to the altered phenotypes, which are often linked to a disease. The distinct interaction profiles caused by genetic variants could provide a more accurate link between genotype and phenotype [17]. We then formulate and test two hypotheses. The first hypothesis states that the phenotypes caused by pathogenic SNVs should be more diverse than the phenotypes caused by frameshift mutations. Our second hypothesis is that the average phenotype similarity score between a pair of a pathogenic SNV and a pathogenic frameshift mutation will be higher than the corresponding similarities between the pairs of pathogenic SNVs.

Each of the two hypotheses is statistically tested with the disease phenotype similarity identified for each pair of mutations [24]. The disease phenotype dissimilarity spans 5,080 diseases in OMIM. The similarity between OMIM records is calculated by comparing the feature vectors, in which an entry represents a MeSH concept. For this work, we annotate only those pairs of disease genes where each gene has at least one pathogenic nsSNV and at least one pathogenic frameshift mutation. For each disease gene, we define the average disease similarity between all nsSNVs on this gene (*S*_1_), between all frameshift mutations associated with the gene (*S*_2_), and between each nsSNV and each frameshift mutation from this gene (*S*_3_) as following:

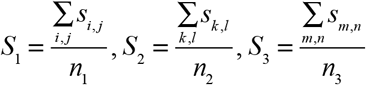

where *s_i,j_* is the similarity score between the phenotype corresponding to SNV *i* and SNV *j* and *n*_1_ is the number of total SNV pairs for the gene; *S*_2_ is defined in the same way for frameshift mutation *k* and frameshift mutation *l*, as well as *S*_3_ is defined for an SNV *m* and a frameshift mutation *n*. To verify each hypothesis, we compare the average similarities defined above, and the statistical significance is calculated using a Mann–Whitney test.

### Examination of pathogenic SNVs in a structurally resolved PPI network

When studying molecular networks centered around a complex disease, the proteins implicated in the disease are often treated as mere network nodes. It has been suggested that adding structural details about the mutations and the corresponding proteins could help in understanding the mutations’ roles in the complex disease. We then examine the distribution of SNVs in a structurally resolved PPI network, INstruct [20]. INstruct is a database of high quality, structurally resolved protein-protein interactions for human. The database includes high-quality binary PPI data and the structural information about the PPI complex at the atomic or near-atomic resolution level derived from the experimental data using a tested interaction interface inference method [20]. For each PPI from INstruct, the PPI interface and the binding sites of each interacting protein have been structurally characterized.

Combining the domain information collected from Uniprot and the PPI interface information collected from INstruct, we divide a disease-related protein into the following three regions: “interface domain”, “non-interface domain” and “non-domain”. If the distributions of pathogenic mutations are not influenced by the domain architecture of the protein, then one should expect for the numbers of SNVs across the three regions to correlate with the lengths of these three regions. Then, the odds ratio (*OR*) for pathogenic SNVs on each of these three regions is calculated. The odds ratio is a statistical measure of association between an exposure and an outcome defined as:

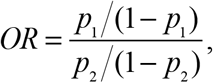

where *p*_1_ is the number of observed mutations in each region across all proteins divided by the total number of mutations and *p*_2_ is the total sequence length of each region across all proteins divided by the length of all proteins combined.

### Functional annotation of SNV’s effect on PPI using SNP-IN tool

Determining whether an nsSNV disrupts or preserves a PPI is a challenging task. We have previously formulated this task as a classification problem, and developed a computational method, SNP-IN tool (non-synonymous SNP INteraction effect predictor tool) [14]. SNP-IN tool predicts the effects of nsSNVs on PPIs, given the interaction’s experimental structure or accurate comparative model. There are three classes of edgetic effects predicted by the SNP-IN tool: beneficial, neutral, and detrimental. The effects are assigned based on the difference between the binding free energies of the mutant and wild-type complexes (*ΔΔG*). The beneficial, neutral, or detrimental types of mutations are then determined by applying two previously established thresholds to *ΔΔG*[51, 52]:

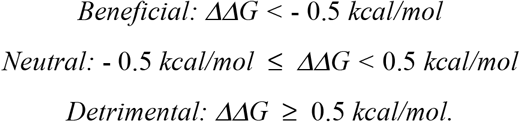

To apply SNP-IN tool, we first structurally characterize, when possible, each PPI from one of the two interaction networks in which a disease protein is involved. If a PPI already has a native structure in PDB, we extract the interaction structure directly using the recently launched INstruct database [20]. Specifically, we identify an interacting chain pair for each PPI in the corresponding PDB file, to make sure that the two chains physically interact. During the process, 3did database [53] is utilized, which maintains the information about the two interacting domains with physical interfaces. If a PPI does not have a structure in PDB, two options are explored. First, if a structural template for such interaction (*i.e.* a homologous protein complex) exists, a comparative model of this interaction can be obtained. Alternatively, if one cannot structurally resolve the full-length PPI, one can try to model only the domain-domain interaction on which the mutation can be mapped to. Homology modeling is done through Interactome3D [25], a web service for structural modeling of PPI network. When modeling a PPI involving either the full-length proteins or partial, domain-domain, structures, the template with the highest sequence identity with respect to the target sequences is selected. Finally, for each PPI that is structurally resolved, the mutated residue is mapped to the protein structure by indicating the position of the mutated residue in the PDB file of the modeled PPI.

### Network cumulative damage analysis

We intend to quantify the cumulative damage effect of a group of pathogenic SNVs on a PPI network associated with a certain disease. To do so, we adapt the methodology from the network robustness theory. The first step of our cumulative damage analysis is to define the “attacking strategy” for the genetic variants. In the traditional network robustness analysis [54], a simple way to perturb a network is to randomly remove its nodes, which we refer to as the “random failure”. Another way is to remove nodes in order of their degrees, from the highest to the lowest, which we call the “malicious attack”. However, these two strategies can be viewed as two extremes. Neither of the two strategies can realistically model the damage caused by the disease genes and pathogenic mutations disrupting the PPIs, and hence these strategies cannot characterize the biological network rewiring behavior. In real networks, the failure or attack could also occur on the edges. In our case, disrupted interactions caused by pathogenic SNV are more likely to fall into this category. The pathogenic mutations would only disrupt limited number of the interactions. It suggests us that an “edge-based” attack strategy might be more suitable to characterize the network rewiring behavior. On the contrary, since frameshift mutations usually result in the polypeptide abnormally short or abnormally long, and the final product will most likely not be functional. They more conform to the complete removal of the nodes. We also included frameshift mutations in this analysis as a comparison. We tried different “attack strategies” (both node-based and edge-based) to find out how pathogenic SNVs perturb the network. For node-based attach strategy, we pick the node degree to guide the node removal process, as previous work reports that degree centrality is proven to be superior to other centrality measures at exposing the vulnerability under malicious attacks and random failures [55]. For edge-based attack strategy, a link-robustness concept based on the highest edge-betweenness attack was recently proposed [19]. This concept captures the network behavior for any fraction of link removal. In this work, we focus on the disease-associated proteins and PPIs disrupted by the pathogenic SNVs according to the SNP-IN tool annotation. And we remove the corresponding edges in the PPI network based on their betweenness centrality.

After we select the “attack strategy”, a quantitative metric to measure the cumulative damage caused by the pathogenic SNVs is defined. One of the key aspects of studying the robustness of a physical networked system against the failure of their component parts is to understand how the size of the largest component changes as nodes and/or edges are removed from the network. If the size of the largest component shrinks sufficiently after the failure, when compared to the original size of the network, then it is reasonable to assume that the networked system is unlikely to function [18]. For an initial network of size *N* with a largest component *S*_0_, removing a fraction of the nodes or edges according to some specified procedure described above would result in a new network, in which the largest component would be *S*_1_. The key quantity that we will study here is the size of *S*_1_ relative to the initial size of the network: *|S*_1_*|/ N*, where *|S*_1_*|* denotes the number of vertices in *S*_1_. Given a set of pathogenic mutations, the cumulative damage to the network caused by the mutations is then defined as: *(|S*_0_*|-|S*_1_*|)/N*. Once a suitable centrality measure and the attack strategy have been fixed, we can compute *|S*_1_*|/N* as a function of the fraction for removed nodes/edges in decreasing order of that centrality measure to characterize the network rewiring behaviors.

### Correlation between disruptive mutations and decreased survival in cancer patients

We next study the relationship between the mutations predicted as disruptive and survival in the cancer patients. To do so we first assemble a list of well-known cancer genes. Specifically, a high-confidence collection of 869 cancer genes is defined as a union of genes in the Cancer Gene Census [42] and recent literature [43]. The Cancer Gene Census catalogs the genes for which mutations have been found implicated in cancer. The recently published dataset [43] was derived using computational methods and includes a list of 291 high-confidence cancer driver genes from 3,205 tumors and 12 different cancer types. Thus, the two datasets are complementary, and we consider their union to be our final dataset. To explore the functional and clinical significance of disruptive mutations on these cancer drivers, we curate the genomic and clinical data for the cancer patients from ICGC. Somatic mutations from several cancer genomics projects are downloaded from the International Cancer Genome Consortium (ICGC) data portal [56]. A subset of mutations mapped to the human genome build 37 are annotated with ANNOVAR [44]. We discard all non-coding and silent mutations, short insertions and deletions, and retain only non-synonymous, missense SNVs.

Finally, we study somatic mutations occurring on the above set of the cancer drivers with high mutation frequency. Specifically, we consider the cancer driver genes with the mutation rate higher than the background mutation rates plus the standard deviation. For those pathogenic mutations, we annotate their effects on the corresponding protein-protein interactions to investigate whether the rewiring of the PPI network would play a role in cancer. Then, based on the mutation annotation results, we divide the patients into two groups: the cancer patients with mutations potentially affecting the interactions and cancer patients without such mutations. Because SNP-IN tool can only be applied to a limited number of these mutations, the obtained groups using SNP-IN tool based annotation are limited in size, preventing application of statistical tests. Therefore, we resort to a more general annotation of somatic mutations by using the information stored in INstruct. More specifically, the interaction interface data for each cancer driver, when available, is extracted from INstruct database, and we check whether the mutation is located in the interface. Lastly, given the annotation results, the comparison of the survival distributions of two groups is performed using the log-rank test [57]. The log-rank test is among the most popular methods for comparing the survival of groups. To do so, the method computes the observed and expected numbers of events in one group at each observed event time, and then obtains the overall summary across all event times as a hazard ratio.

## Acknowledgements

D.K. acknowledges funding from the US National Science Foundation (award DBI-1458267).

## Author contributions statement

H.C. and D.K. conceived the experiment(s), H.C. conducted the experiment(s), H.C. and N.Z. performed the functional annotation of mutations, H.C. and D.K. analyzed the results. H.C. and D.K. wrote the manuscript. All authors reviewed the manuscript.

